# Re-evaluating rhythmic attentional switching: Spurious oscillations from shuffling-in-time

**DOI:** 10.1101/2021.05.07.443101

**Authors:** Geoffrey Brookshire

## Abstract

How does attention help to focus perceptual processing on the important parts of a visual scene? Although the neural and perceptual effects of attention were traditionally assumed to be sustained over time, the field is converging on a dramatically different view: that covert attention rhythmically switches between objects at 3-8 Hz. Here I demonstrate that ubiquitous analyses in this literature conflate rhythmic oscillations with aperiodic temporal structure. Using computational simulations, I show that the behavioral oscillations reported in this literature could reflect aperiodic dynamics in attention, rather than periodic rhythms. I then propose two analyses (one novel and one widely used in climate science) that discriminate between periodic and aperiodic structure in behavioral time-series. Finally, I apply these alternative analyses to published data-sets, and find no evidence for rhythms in attentional switching after accounting for aperiodic temporal structure. Attention shows rich temporal structure. The techniques presented here will help to clarify the periodic and aperiodic dynamics of perception and cognition.

## Introduction

Our senses provide us with a vast amount of simultaneous information. Attention helps to focus perceptual processing on the important parts of a scene, boosting both perceptual sensitivity and neural responses to a stimulus [1, 2]. The effects of attention were traditionally assumed to be sustained over short timescales; as long as someone holds their attention on a stimulus, processing of that stimulus is boosted. However, the field is now converging on a dramatically different view: that covert attention rhythmically switches between objects at 3-8 Hz [3–6]. Here I show that ubiquitous analyses in this literature conflate periodic oscillations with aperiodic temporal structure, leading to drastically inflated rates of statistical false positives. I then present two alternative analyses that can distinguish between periodic and aperiodic temporal structure in behavior while controlling the rate of false positives.

A growing behavioral literature argues that the focus of attention moves rhythmically between stimuli several times per second [7–27]. The experiments in this literature use a variety of stimuli, but are built on a shared core design. In these studies, participants monitor two peripheral stimuli for a faint target. After a short delay, a cuing stimulus flashes on the screen. This cue has been hypothesized to reset the phase of low-frequency neural oscillations, and serves to draw attention to one of the two peripheral stimuli. After a second variable delay, a faint target flashes on one of the peripheral stimuli. By averaging target-detection accuracy at each cue-to-target delay, these studies create a time-course of attention toward the cued and uncued locations. To identify rhythms in attentional switching, amplitude spectra are then computed for these behavioral time-courses. Peaks in the spectra are interpreted as evidence that attention moves rhythmically around the perceptual scene.

Studies of rhythmic attentional switching have used a wide range of different stimuli and dependent variables. For example, some studies examine visual attention to different spatial locations [7, 8, 10], whereas others focus on feature-based attention [21], global-local processing [28], or on auditory attention [15]. These studies have reported rhythms in detection accuracy [7, 8], reaction times [9], binocular rivalry [18], cue validity effects [26], and in sensitivity and criterion metrics from signal detection theory [15]. Most studies reset ongoing dynamics with a cue stimulus, but some rely on participant-initiated actions [12]. Although prominent theories focus on rhythms around 4-8 Hz [4–6], studies in this literature have reported behavioral rhythms as low as 2.5 Hz [16] and as high as 20 Hz [28]. This large and diverse literature is widely interpreted as converging evidence for robust rhythms in attentional switching.

Here I demonstrate that the findings in this literature can be accounted for by attentional switching that is entirely non-oscillatory. Using computational simulations, I demonstrate that the spectral analyses used in this literature are sensitive not only to periodic rhythms, but also to aperiodic temporal structure. I present two alternative methods that discriminate between periodic and aperiodic structure, control the rate of false positives, and recover true oscillations in behavior.

## Results

### Identifying oscillations by shuffling in time

Does attention move rhythmically between different objects? A large number of studies have addressed this question by searching for oscillations in densely-sampled behavioral time-series. After converting this time-series to the frequency domain, oscillations appear as peaks in the amplitude spectrum. To interpret this spectrum, any putative oscillations must be discriminated from the background noise. How can we test whether a peak in the spectrum is significantly greater than the background noise?

Studies in this literature test for statistically significant oscillations by performing a randomization procedure that relies on shuffling the data in time. By shuffling in time, this analysis creates a surrogate distribution without any temporal structure, and searches for oscillations against this surrogate distribution. We illustrate the basic procedure using details from an early influential study [7] (LF2012; Fig. 1). First, accuracy is computed at each time-point, yielding a densely-sampled behavioral time-series. The data are then detrended with a 2nd-order polynomial, multiplied by a Hanning taper, and zero-padded, before computing the amplitude spectrum with a discrete Fourier transform (DFT). To test whether peaks in this spectrum are statistically significant, a randomization test is performed. The time-stamps of the raw behavioral data are shuffled a large number of times, and then the spectra of these time-shuffled data are computed. This results in a surrogate distribution of randomized spectra. For each frequency, the p-value is computed as the proportion of randomized spectra with greater amplitude than the empirical spectrum. P-values are then corrected for multiple comparisons using Bonferroni corrections.

**Figure 1:**
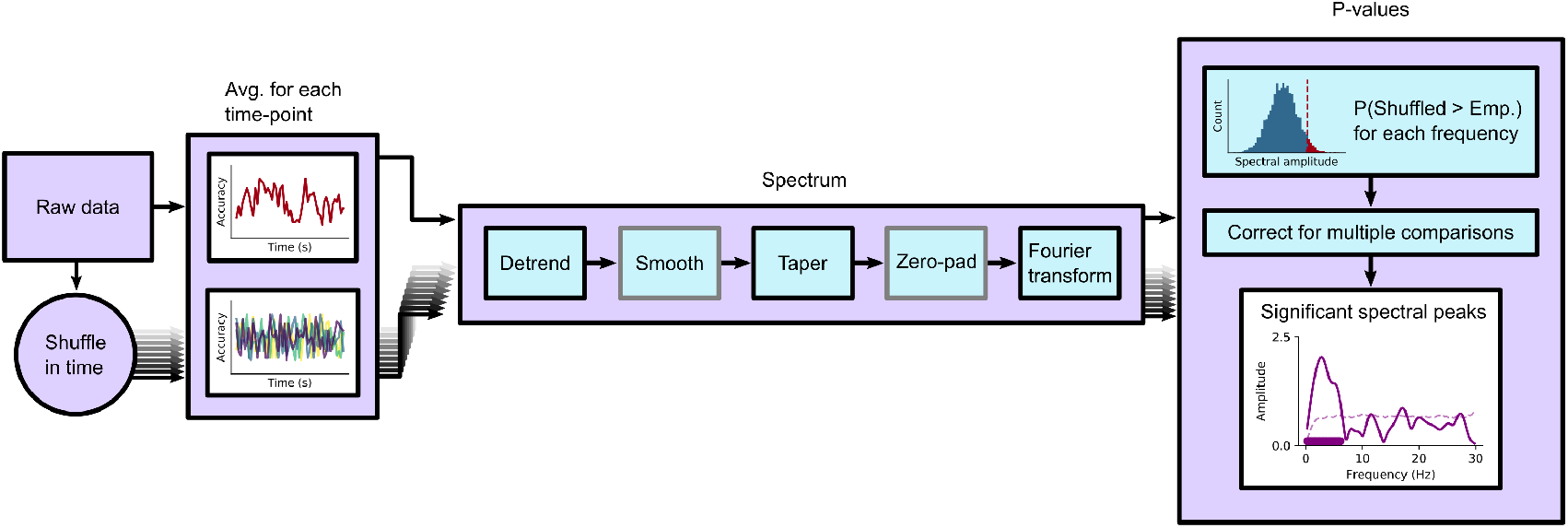
Standard analysis pipeline: identifying behavioral oscillations by shuffling in time. Schematic of the standard analysis pipeline used to test for statistically significant rhythms in behavioral data. The analysis steps shown in grey are not performed in every study. Different researchers sometimes perform some steps in different ways, for example by detrending with a 2nd-order polynomial versus a sliding rectangular window. In the panel labeled ‘Significant spectral peaks’, the solid line indicates the empirical spectrum, the dotted line shows the significance threshold, and the solid bar shows frequencies that were identified as statistically significant.

This shuffling-in-time procedure is widely used to study attentional switching [7–27] as well as other rhythms in perception [28–32]. The details of the analysis pipeline often differ between studies. For example, instead of computing accuracy at each time-point, some studies use different dependent measures, such as average RT [9] or d-prime [15]. Some studies do not zero-pad the data before computing the amplitude spectrum [10], or they correct for multiple comparisons across frequencies using the false discovery rate [8] or by selecting the largest peak in each shuffled spectrum [21]. All of these studies, however, determine statistical significance using a randomization test that shuffles the raw data in time.

### Shuffling in time alters aperiodic temporal structure

Significant spectral peaks from the shuffling-in-time procedure are interpreted as reflecting periodic rhythms in attention. By shuffling the data in time, however, these studies test the null hypothesis that the behavioral data have no structure in time. These tests do not, therefore, provide unique evidence for oscillations in behavior. Instead, they provide evidence for any kind of structure in time. Randomization tests which shuffle the data in time reflect at least two varieties of non-oscillatory structure: aperiodic autocorrelation and consistency over trials.

First, shuffling in time destroys aperiodic temporal structure due to autocorrelation. Autocorrelation refers to correlations between a signal and lagged copies of itself. For periodic signals, the autocorrelation function has regularly-spaced bumps, showing that the signal is positively correlated with itself at those periodic lags. For aperiodic signals, however, a different pattern emerges. For example, in a random walk, the signal at time *t* is strongly correlated with itself at time *t* – 1, but weakly correlated with itself at more distant times. As a consequence, the autocorrelation function smoothly drops down to zero with increasing lags. When data are simulated using a noisy random walk (an autoregressive model with a single positive coefficient [AR(1)]; Fig. 2a), the autocorrelation function slowly drops to zero (Fig. 2c). After shuffling in time (Fig. 2b), however, the autocorrelation is approximately zero at all non-zero time-lags (Fig. 2c). This difference in the autocorrelation functions also appears in the amplitude spectra. When those data are preprocessed using detrending and other common analysis steps, an aperiodic autocorrelated signal often shows a single spectral peak, even though the data reflect a purely aperiodic process (Fig. 2d).

**Figure 2:**
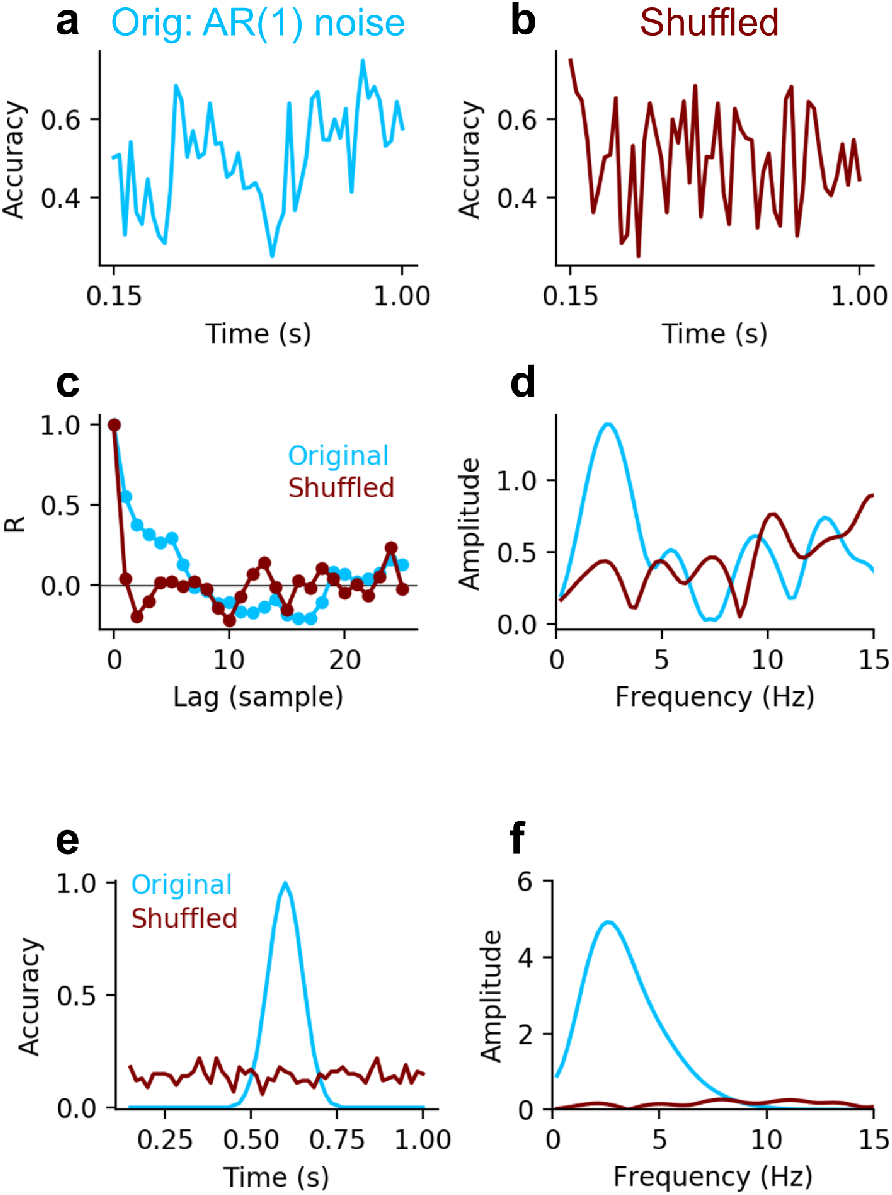
Shuffling in time alters aperiodic temporal structure. **(a-d)** Autocorrelation is destroyed by shuffling in time. **(a)** Simulated behavioral data generated with an aperiodic AR(1) function (*β* = 0.5). **(b)** The same time-series after being shuffled. **(c)** Autocorrelation of the original AR(1) and shuffled timeseries. **(d)** Amplitude spectra of the original AR(1) and shuffled time-series, computed using the pipeline from LF2012. Shuffling in time introduces an apparent peak in the spectrum of the AR(1) process, even though the autocorrelation function does not suggest that the signal is periodic. **(e-f)** Consistency across trials is destroyed by shuffling in time. **(e)** A simulated behavioral time-series with a single peak in accuracy, and accuracy computed after shuffling trials. **(f)** Amplitude spectra of the original and shuffled data, computed using the pipeline from LF2012.

Second, shuffling in time destroys aperiodic consistency across trials. This is illustrated by simulating a behavioral time-course with a single peak in accuracy (Fig. 2e). After shuffling the time-points between trials, that single peak in accuracy is spread out over all time-points. In the amplitude spectra, this results in decreased amplitude at a wide range of frequencies, with a single peak that depends on the shape of the original aperiodic signal (Fig. 2f).

In summary, shuffling in time tests the null hypothesis that a time-series contains no temporal structure whatsoever. However, this method cannot distinguish between periodic rhythms and aperiodic temporal structure.

### Distinguishing between periodic and aperiodic temporal structure

Shuffling in time tests the null hypothesis that a time-series has no temporal structure of any kind. Here, we outline two procedures that can discriminate between periodic and aperiodic temporal structure in behavioral time-series. The first procedure is a novel randomization method based on autoregressive models. The second procedure, based on harmonic analysis with the multi-taper method, is commonly used to identify rhythms in climate science [33].

The novel method is based on randomization, but creates the surrogate distribution using an auto-regressive model instead of by shuffling in time. Let’s call this the ‘AR surrogate’ analysis (Fig. 3). First, the empirical time-series is obtained by computing accuracy (or some other aggregated measure) at each time-point. Next, an autoregressive model with one positive coefficient [AR(1)] is fit to this time-series. This AR(1) model captures the aperiodic structure - but not the periodic structure - of the time-series [33]. This fitted AR(1) model is then used to generate a large surrogate distribution of time-courses that preserve the aperiodic structure of the original signal. For the empirical and surrogate time-courses, the amplitude spectrum is obtained using a discrete Fourier transform. P-values are computed for each frequency as the proportion of surrogate spectra with greater amplitude than the empirical spectrum. Finally, a cluster-based permutation test [34] is used to correct for multiple comparisons across frequencies.

**Figure 3:**
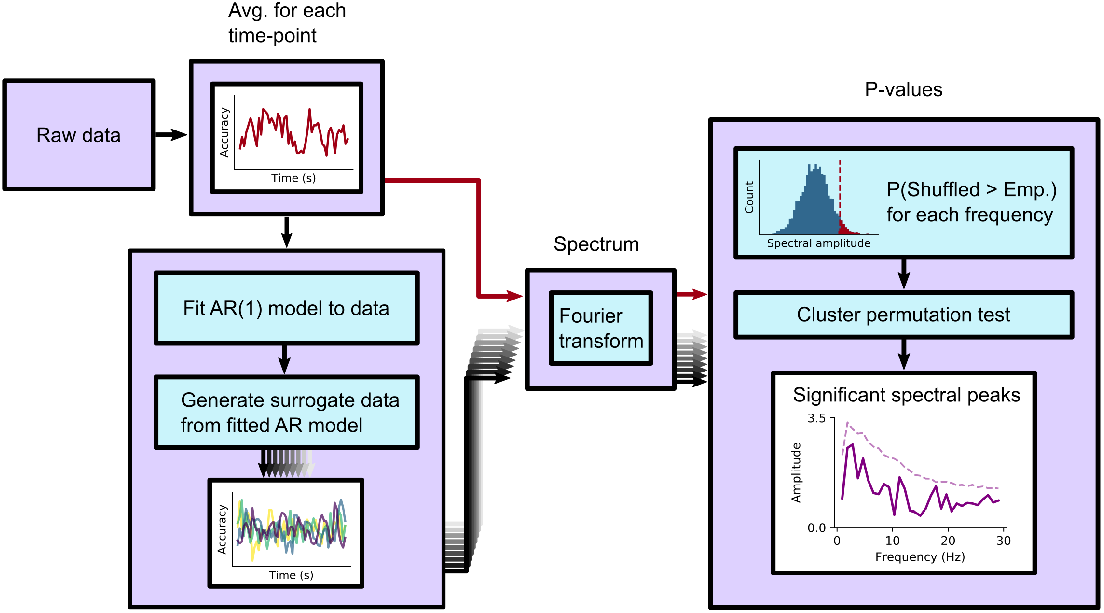
AR surrogate analysis for identifying oscillations in behavior. Procedure to test for behavioral oscillations by generating surrogate data with a fitted AR(1) model.

For the second analysis method, we use a procedure that is widespread in climate science [33]. This method, called the ‘robust estimate’, was developed to identify oscillations in autocorrelated background noise in geological time-series. Because human behavior is also autocorrelated [35, 36], this method may help distinguish behavioral rhythms from aperiodic background activity. This method uses multi-taper spectral analysis to compute the power spectrum of the signal, removes narrow-band peaks with median-smoothing, and then makes a robust estimate of the background noise by fitting an analytic AR(1) noise spectrum. Finally, statistical significance is computed by comparing the empirical spectrum to the AR(1) background fit [33].

### Shuffling in time inflates the rate of false positives

Aperiodic temporal structure may appear as peaks in the amplitude spectrum when analyzed by shuffling in time. To test how the different analysis methods reflect periodic and aperiodic structure, I simulated behavioral experiments of attentional switching. Experiments were simulated following the methods and analyses of two foundational studies in this literature: Landau and Fries, 2012 (LF2012); and Fiebelkorn, Saalmann, and Kastner, 2013 (FSK2013). To explore the novel AR surrogate and robust estimate analyses, I simulated experiments following the behavioral paradigm in LF2012.

To examine how aperiodic structure can lead to false positive results, I simulated four types of temporal structure. First, I simulated experiments in which every trial was independently and randomly determined to be a hit or a miss (‘fully random’). These experiments had no temporal structure at all, and act as a baseline for each analysis method. All four analysis methods yield false positives around or below the expected rate of *α* = 0.05 (Fig. 4a; Table 1), indicating that they avoid false positives when testing against the null hypothesis that behavior does not contain any temporal structure.

**Figure 4:**
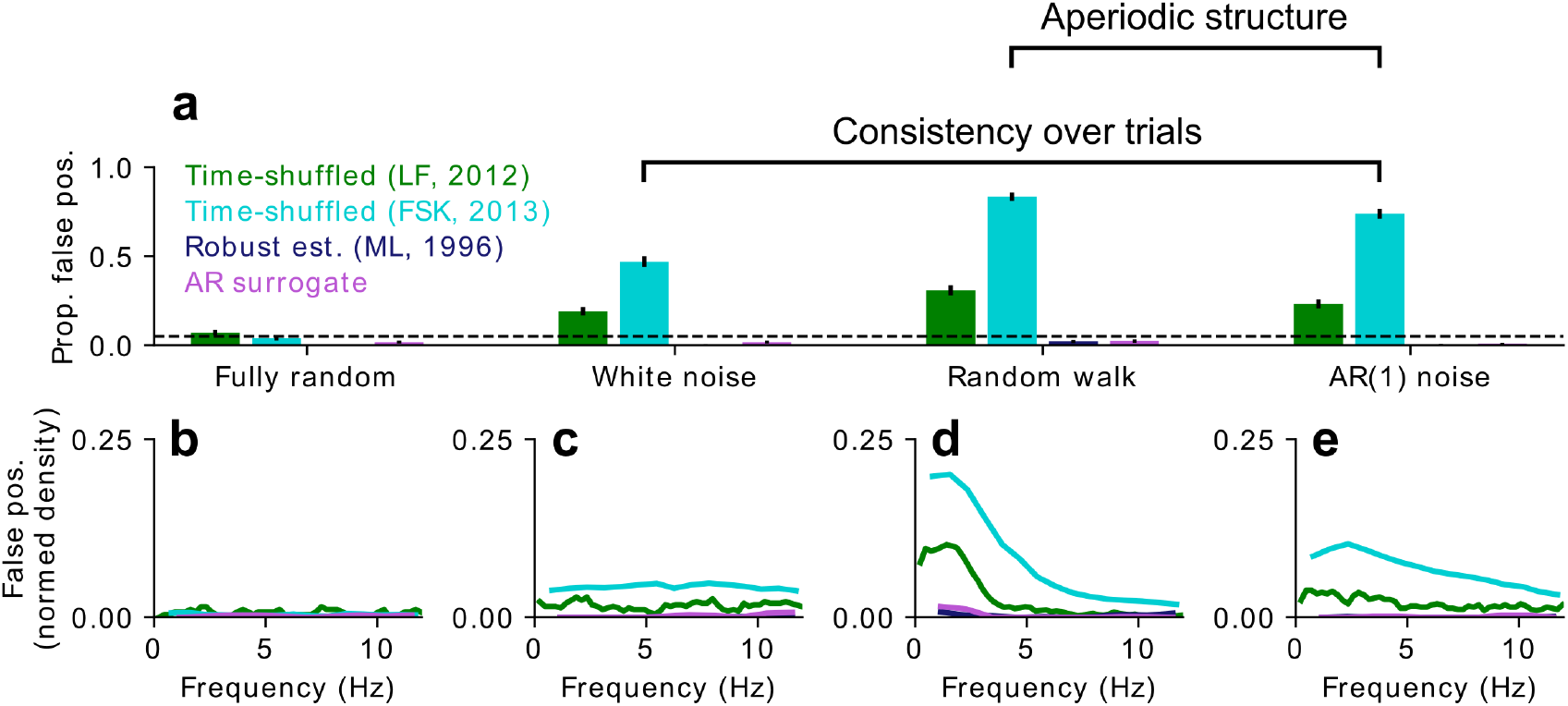
False positive oscillations from aperiodic signals. **(a)** Proportion of false positive results in simulated experiments. All noise processes except fully random noise include consistency over trials, and non-zero autocorrelations are present in both random walk and AR(1) noise. Error bars show 95% confidence intervals. Each bar includes data from 1000 simulated experiments. The dotted line shows the expected rate of false positives (*α* = 0.05). **(b-e)** For each type of aperiodic noise, the rate of false positives by frequency for each analysis method. **(b)** Fully random. **(c)** White noise. **(d)** Random walk. **(e)** AR(1) noise. Colors as in **(a)**. False positives have been normed by the number of frequency bins in each analysis method, to allow for comparisons across methods, and scaled to reflect the total number of false positives in each method. The exact frequency bins differ across methods due to differences in how each method computes the amplitude spectrum.

**Table 1:** Proportion of false positives for each analysis method and noise type. P-values (in parentheses) reflect the difference from the expected false-positive rate of 0.05 (one-tailed binomial tests). Proportions that are significantly greater than .05 are shown in bold. *N* = 1000 simulated experiments in each cell.

|  | LF2012 | FSK2013 | Robust est. | AR surr. |
| --- | --- | --- | --- | --- |
| Fully random | **0.07** (.002) | 0.04 (.95) | 0.000 ($\approx 1$) | 0.018 ($\approx 1$) |
| White noise | **0.19** ($10^{-56}$) | **0.47** ($\approx 0$) | 0.002 ($\approx 1$) | 0.018 ($\approx 1$) |
| Random walk | **0.31** ($10^{-150}$) | **0.84** ($\approx 0$) | 0.022 ($\approx 1$) | 0.024 ($\approx 1$) |
| AR(1) noise | **0.23** ($10^{-86}$) | **0.74** ($\approx 0$) | 0.003 ($\approx 1$) | 0.009 ($\approx 1$) |

To investigate how these methods perform with behavior that is consistent across trials, I simulated experiments in which the response in each trial was randomly determined according to an idealized accuracy function. This function specified the time-course of accuracy that would be obtained with an infinite number of trials. Experiments were simulated for three types of idealized accuracy time-courses. In ‘white noise’ simulations, idealized accuracy time-courses were generated with a random Gaussian process; these simulations included consistency over time but no other temporal structure. In ‘random walk’ simulations, idealized accuracy time-courses were generated with a random walk, including both consistency over trials and aperiodic temporal structure. Random walks, also called Brownian noise and 1/*f*^2^ noise, form the basis for various models in neuroscience [37] and psychology, including drift-diffusion models of decision-making [38, 39]. Finally, in ‘AR(1) noise’ simulations, idealized accuracy time-courses were generated with an autoregressive process with one coefficient (*β* = 0.5); these simulations include both consistency over trials and aperiodic temporal structure.

Analyses based on shuffling in time (LF2012 and FSK2013) showed strongly inflated rates of false positives when the simulated data were consistent over trials (Fig. 4a, ‘white noise’). False positive rates were higher still when the simulated data included autocorrelational structure (Fig. 4a, ‘random walk’ and ‘AR(1) noise’). In contrast, the analyses that do not rely on shuffling in time (AR surrogate and robust est.) had low rates of false positives for all noise types. False positives when shuffling in time appeared at a wide range of frequencies, with the most common frequencies depending on the process used to generate the noise. In white noise simulations, false positives were evenly distributed across frequencies (Fig. 4c), reflecting the flat spectrum of white noise. In random walk simulations, false positives were biased toward lower frequencies, with a dip at very low frequencies (Fig. 4d) due to detrending before computing the Fourier transform. False positives in AR(1) noise simulations were intermediate between the prior two, with slightly higher rates of false positives at lower frequencies (Fig. 4e).

In summary, shuffling in time leads us to conclude that behavior is rhythmic even when behavior is generated using a purely aperiodic process. Shuffling in time often generates plausible-seeming amplitude spectra with significant peaks at theoretically interesting frequencies (Fig. 5a-d). In contrast, the two alternative analysis methods (AR surrogate and robust est.) control the rate of false positives for aperiodic processes.

**Figure 5:**
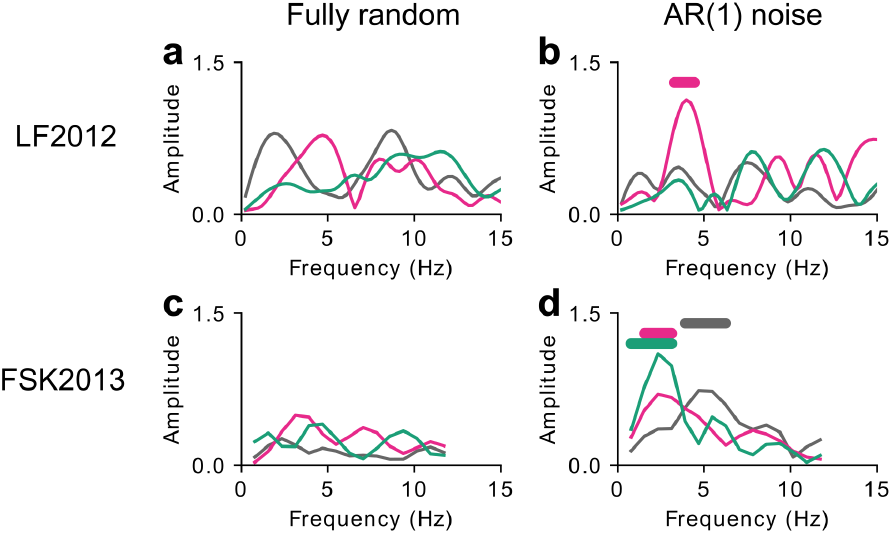
Examples of false positive rhythms from aperiodic noise. Each panel illustrates three randomly-selected simulated experiments (shown in different colors). Bars at the top of the panels show frequencies that were identified in each experiment as significant oscillatory components. **(a, c)** Experiments generated with fully random data. **(b, d)** Experiments generated with AR(1) noise that was consistent over trials. **(a, b)** Data analyzed using the procedure from LF2012. **(c, d)** Data analyzed using the procedure from FSK2013.

### Alternative methods recover true oscillations

We can avoid false positive results by using analysis methods that do not rely on shuffling in time (AR surrogate and robust est. methods). Can these alternative methods also recover true oscillations in behavior? To find out, experiments were simulated by combining random walk noise with sinusoidal modulation. These simulations examine the rate of statistically significant results as a function of the frequency and amplitude of behavioral oscillations. Amplitude was coded as the difference in accuracy between peaks and troughs of the idealized oscillation. For example, with amplitude 0.2, accuracy oscillates between 0.4 and 0.6.

These analyses report the proportion of simulated experiments that successfully recovered the behavioral oscillation. This measure is akin to an estimate of experimental power, assuming behavioral data includes random-walk background noise. Both the AR surrogate method and the robust est. method successfully recover true oscillations in simulated behavior. These methods most effectively recover oscillations at higher frequencies and higher amplitudes (Fig. 6a-b), and recover few behavioral rhythms below 4 Hz or below an amplitude of 0.3. The AR surrogate method outperformed the robust est. method above 5 Hz, but the robust est. method performed better at lower frequencies (Fig. 6c). Neither method reliably identified oscillations at 2 Hz; at frequencies that are close to the frequency resolution of the spectra, the AR surrogate method returns very few positive results (Fig. 6a), and the frequencies identified by the robust est. method were often inaccurate by several Hz (Fig. 6e). At higher frequencies, however, the peaks identified by both methods are highly accurate and precise (Fig. 6d-f). Averaging over frequencies of 3 Hz and above, and amplitudes of 0.2 and above, the frequency of recovered oscillations was the most accurate for the AR surrogate method. Both the AR surrogate and robust est. methods were more accurate than methods relying on shuffling in time (Fig. 6g-h, Table 2).

**Figure 6:**
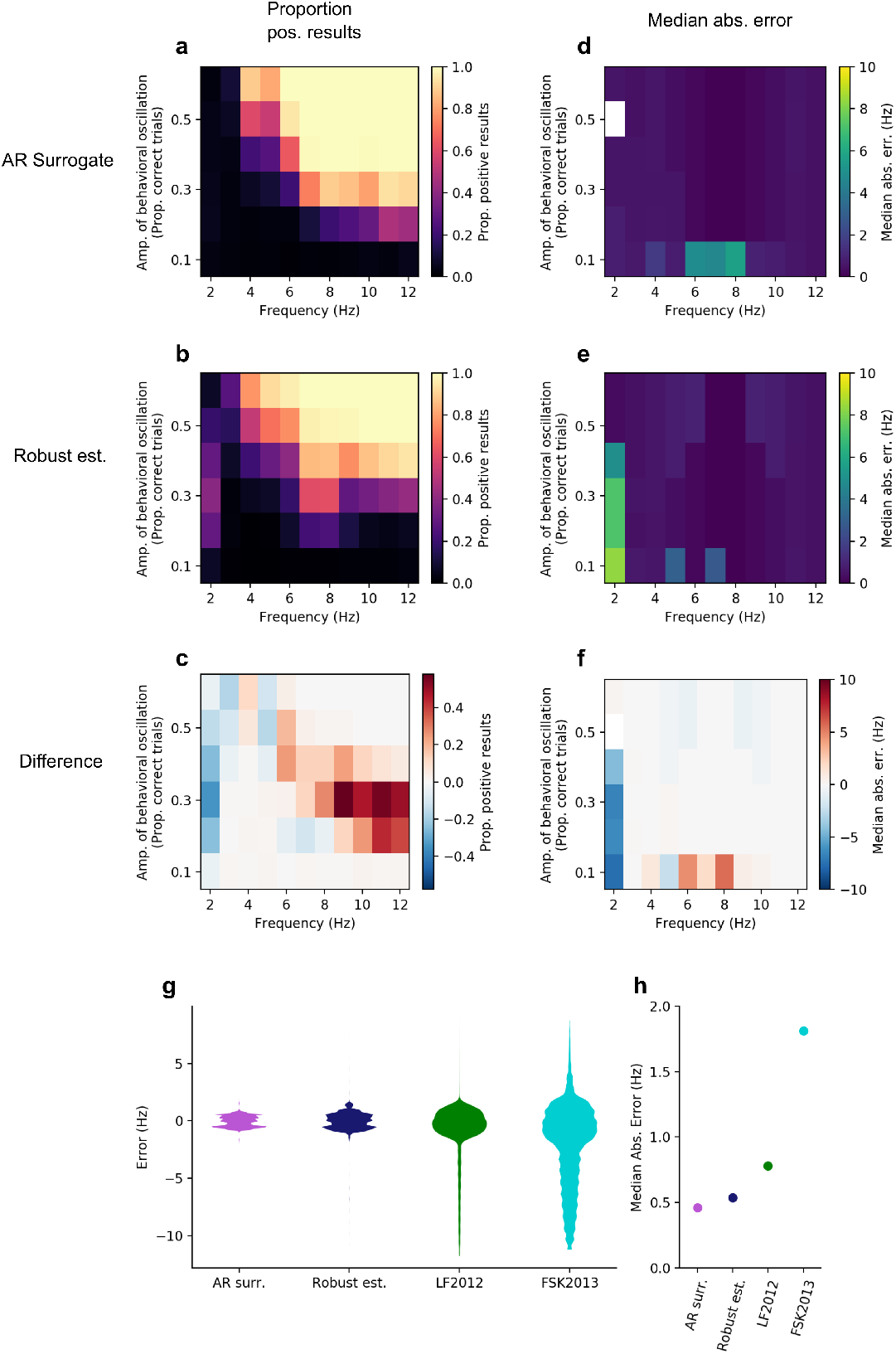
AR surrogate and robust est. methods recover oscillations in simulated behavior. **(a)** Proportion of simulated experiments that correctly recover oscillations when analyzed with the AR surrogate method, plotted as a function of the frequency and amplitude of behavioral oscillations. Color corresponds to the proportion of experiments that found significant oscillations in behavior at *p* < .05. Each cell shows the results from 1000 simulated experiments. **(b)** As in **(a)**, but analyzed with the robust est. method. **(c)** Difference in proportion of recovered oscillations between the AR-surrogate and robust est. methods. Positive values correspond to higher experimental power for the AR surrogate method, and negative values to higher experimental power for the robust est. method. **(d)** The median absolute error of frequencies identified as significant using the AR surrogate method. **(e)** As in **(d)**, but using the robust est. method **(f)** Difference in median absolute error between the AR surrogate and robust est. methods. Positive values correspond to more accurate frequency estimates using the robust est. method, and negative values to more accurate frequency estimates using the AR surrogate method. White cells in **(d-f)** correspond to simulations with no statistically significant results. **(g)** Error of the statistically-significant frequencies for each method. Violin plots depict the full distribution of errors. **(h)** The median of the absolute value of the errors for each analysis method. Results in **(g-h)** include simulated frequencies ≥ 3 Hz, amplitudes ≥ 0.2.

**Table 2:** Statistical tests on the precision of recovered oscillations. Each row shows the results of a linear regression on the median absolute error (MAE) of the reconstructed frequency of true oscillations. Regressions compared each pair of analysis methods (e.g. AR surrogate vs. Robust est.), and included terms to control for the amplitude and frequency of the simulated oscillation, with degrees of freedom (3, 96). MAE was entered into the regression for each frequency and amplitude of the simulated oscillations, and considered simulated frequencies ≥ 3 Hz, and amplitudes ≥ 0.2.

| Comparison | Est. | 95% CI | t | p-value |
| --- | --- | --- | --- | --- |
| AR surr., Robust est. | 0.09 | -0.01, 0.18 | 1.88 | 0.063 |
| AR surr., LF2012 | 0.41 | 0.35, 0.47 | 12.85 | $1.4 \times 10^{-22}$ |
| AR surr., FSK2013 | 1.88 | 1.54, 2.21 | 11.06 | $7.8 \times 10^{-19}$ |
| Robust est., LF2012 | -0.32 | -0.40, -0.25 | -8.87 | $4.1 \times 10^{-14}$ |
| Robust est., FSK2013 | -1.79 | -2.13, -1.45 | -10.45 | $1.6 \times 10^{-17}$ |
| LF2012, FSK2013 | -1.47 | -1.80, -1.13 | -8.63 | $1.3 \times 10^{-13}$ |

How do these alternative methods compare to the standard approach of shuffling in time? Shuffling in time could identify a large proportion of true oscillations, but results from these methods would be difficult to interpret if they also produce a large number of false positives. To quantify this trade-off, I computed the ratio of correct positive results to false positives when no oscillation is present (‘detection ratio’). When data were analyzed by shuffling in time, the detection ratio was low (< 3.5) for all frequencies and amplitudes (Fig. 7a-b). The AR surrogate and robust est. analyses, however, showed much stronger detection ratios, especially at high frequencies and amplitudes (Fig. 7c-d). Averaging over frequencies of 3 Hz and above, and amplitudes of 0.2 and above, both of the alternative methods showed substantially higher detection ratios than the methods that use shuffling in time (Fig. 7e, Table 3).

**Figure 7:**
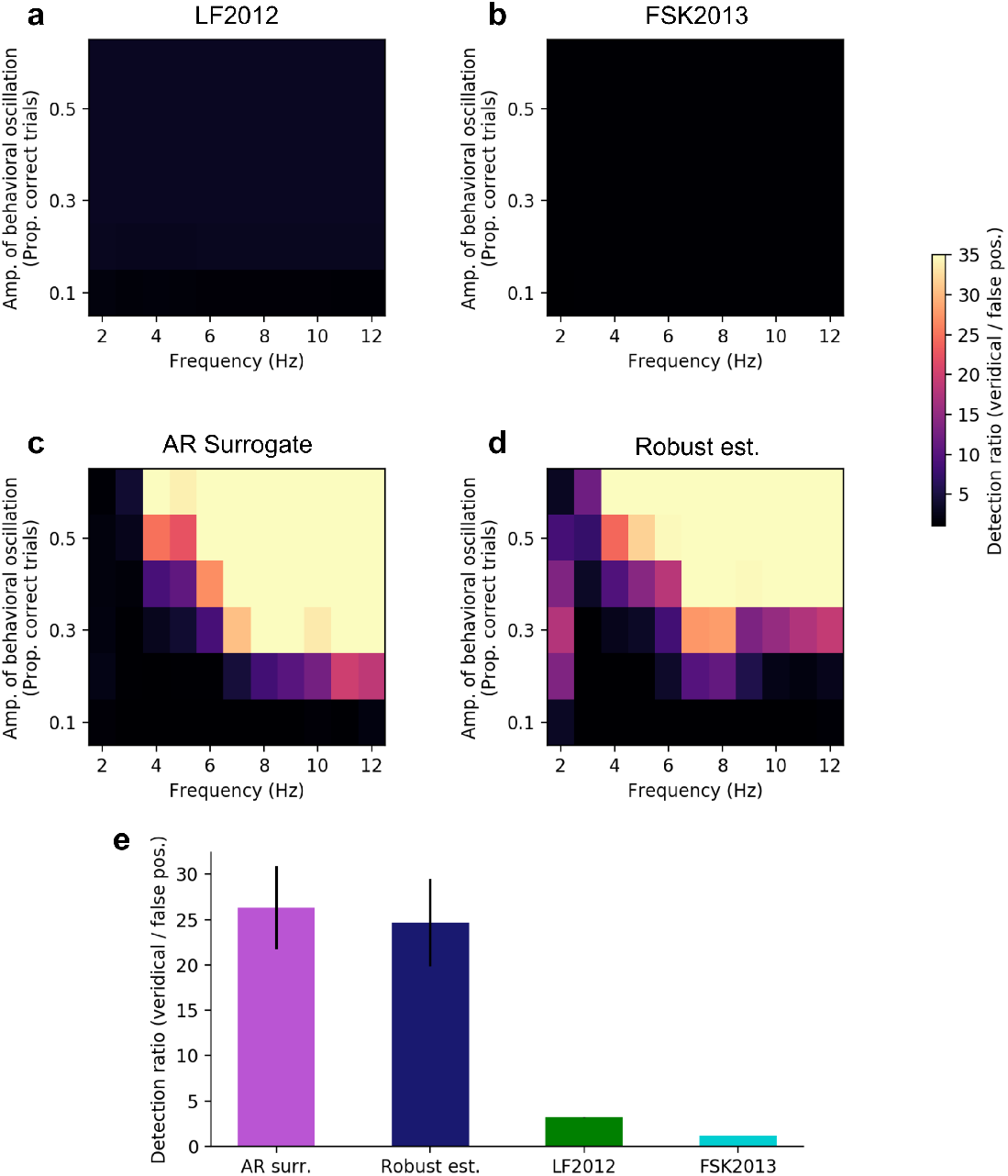
Ratio of correct positive results to false positives. Detection ratio is computed as the proportion of significant results when the simulated data include oscillations, divided by the proportion of significant false positives when the data do not include oscillations. Plotted as a function of the frequency and amplitude of simulated behavioral oscillations, shown separately when analyzed using **(a)** the LF2012 method; **(b)** the FSK2013 method; **(c)** the AR surrogate method; and **(d)** the Robust estimate method. **(e)** Detection ratio for each analysis method, averaged over each cell of the simulations, including simulated frequencies ≥ 3 Hz, amplitudes ≥ 0.2. Error bars show 95% confidence intervals.

**Table 3:** Statistical tests comparing the detection ratios of the different analysis methods. Each row shows the results of a linear regression comparing two analysis methods (e.g. AR surrogate vs. Robust est.). Regressions include terms to control for the amplitude and frequency of the simulated oscillation, with degrees of freedom (3, 96). Detection ratio was entered into the regression for each frequency and amplitude of the simulated oscillations, and included frequencies ≥ 3 Hz, and amplitudes ≥ 0.2.

| Comparison | Est. | 95% CI | t | p-value |
| --- | --- | --- | --- | --- |
| AR surr., Robust est. | -1.64 | -5.08, 1.79 | -0.95 | 0.35 |
| AR surr., LF2012 | -23.12 | -26.84, -19.39 | -12.33 | $1.7 \times 10^{-21}$ |
| AR surr., FSK2013 | -25.12 | -28.85, -21.39 | -13.37 | $1.2 \times 10^{-23}$ |
| Robust est., LF2012 | 21.47 | 17.61, 25.34 | 11.03 | $9.2 \times 10^{-19}$ |
| Robust est., FSK2013 | 23.48 | 19.60, 27.35 | 12.03 | $7.1 \times 10^{-21}$ |
| LF2012, FSK2013 | 2.00 | 1.98, 2.03 | 174.53 | $5.9 \times 10^{-122}$ |

### False-positives in published studies

In principle, shuffling-in-time could cause researchers to find spurious rhythms in non-rhythmic behavior. Alternatively, positive findings in this literature could reflect true attentional rhythms. To distinguish between these possibilities, I re-analyzed behavioral time-courses in publicly-available data [15, 18, 22, 26]. These 4 published studies reported 11 statistically significant behavioral oscillations out of 23 tested timecourses. When re-analyzed using the AR surrogate and robust est. methods, none of these tests reached statistical significance with either analysis method (Fig. 8). This finding suggests that rhythms in behavior may not be as prevalent as the published literature suggests.

**Figure 8:**
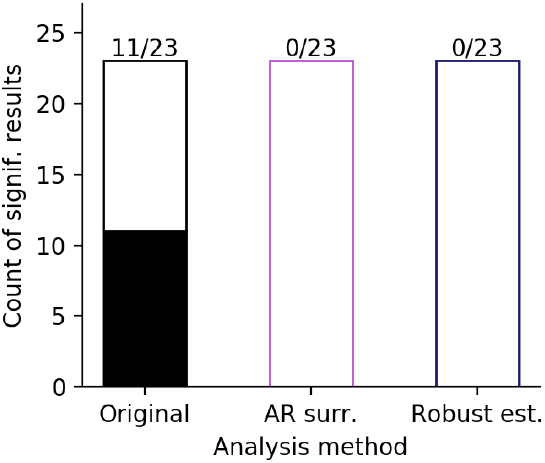
No behavioral oscillations in re-analyses of published data-sets. Re-analysis of publicly-available data from published studies. Each bar shows the number of significant results (filled portion of the bar), and the number of non-significant results (unfilled portion of the bar). The ‘Original’ bar shows the statistical tests reported in the original studies (all based on shuffling in time), and the ‘AR surr.’ and ‘Robust est.’ bars show the results after re-analyzing those data with the alternative analysis methods. Numbers at the top of each bar show (the number of significant results) / (the total number of statistical tests).

## Discussion

What is the temporal structure of attention? Although the field is approaching consensus that attention moves rhythmically, the computational simulations presented here suggest that this conclusion is premature. Shuffling in time tests the null hypothesis that a behavioral time-course shows no temporal structure. This analysis therefore yields positive results whenever the data show any structure in time, regardless of whether that structure is oscillatory. The alternative analyses proposed here (AR surrogate and robust est. methods) test the null hypothesis that the data are autocorrelated but not oscillatory, allowing us to disentangle periodic structure from aperiodic structure. When applied to published data-sets, these alternative analyses do not find any significant oscillations in behavior.

These findings suggest that attention may not switch rhythmically after all. However, the simulations and analyses reported here are consistent with the now-widespread conclusion that attention is non-stationary [7, 8]. These methods will help to clarify the rich temporal structure of attention.

### Anti-phase relationships in behavior

A number of studies have argued for rhythmic attentional switching on the basis of an anti-phase relationship in the spectra of behavior toward the cued and uncued stimuli [7–9, 11, 13, 15]. This anti-phase relationship, they suggest, indicates that attention switches between locations at the frequency of the spectral peak. Although an anti-phase relationship does suggest that attention switches between stimuli, it does not necessarily reflect *rhythmic* switching. For example, inverting a white-noise time-course leads to an anti-phase relationship without any oscillations. Anti-phase dynamics are further evidence of rich temporal dynamics in attention, but they do not uniquely point to periodic rhythms in behavior.

### What null hypothesis does each analysis test?

When we shuffle the data in time, we test the null hypothesis that the data show no structure in time whatsoever. Any structure in time, therefore, can lead to significant results in those tests. For example, shuffling in time gives a positive result if the data show either consistency over trials (e.g. accuracy tends to be lower immediately following a cue stimulus), or autocorrelation (e.g. accuracy at 300 ms is more similar to accuracy at 333 ms than to accuracy at 1000 ms). Shuffling in time, therefore, does not uniquely identify oscillations - instead, it tests whether the data show any kind of (non-specific) temporal structure.

In contrast, the two alternative methods specifically account for non-oscillatory temporal structure. The AR surrogate method fits a model of the data that captures autocorrelational structure, and uses this model to generate a surrogate distribution. This method therefore tests the null hypothesis that the data are autocorrelated but not rhythmic. In other words, the null hypothesis for the AR method is that the data follow a non-oscillatory “red noise” pattern (or equivalently, a noisy random walk or AR(1) process).

The robust est. method uses a different analytic approach, but also tests the null hypothesis that the data follow a non-oscillatory red noise pattern. The robust est. method was designed to test for rhythms in climate data. Oscillations in climate data are embedded in autocorrelated noise, similar to theorized oscillations in behavioral data. In the robust est. method, the data’s spectrum is compared with the best-fitting spectrum of a non-oscillatory autocorrelated process (which has an analytic form); oscillations are identified as points at which the spectrum is significantly higher than the best-fitting autocorrelation spectrum. The robust est. method, therefore, tests the null hypothesis that the data follow a non-oscillatory red noise pattern.

### Do the alternative methods specifically identify oscillations?

Given a strong oscillation in behavior, the AR surrogate and robust est. analyses can positively identify that oscillation. These methods identify oscillations with higher precision (Fig. 6) and higher selectivity (Fig. 7) than the standard approach of shuffling-in-time. The robust est. method has been widely used in climate science for 25 years, helping to identify real oscillations in the presence of autocorrelated noise.

But what can we say about the converse situation: Given a positive statistical result, how confident can we be that an oscillatory process gave rise to that result? In electrophysiological studies, non-rhythmic evoked activity (such as an event-related potential; ERP) can sometimes appear as a brief increase in power at a particular frequency band. Could a similar non-oscillatory process give rise to significant results using the AR surrogate and robust est. methods? In principle, attentional re-orienting could show a pattern analogous to ERPs, in which the cue event is followed by a consistent non-oscillatory response. If that behavioral ‘ERP’ appears as a sequence of 2 to 3 fairly rhythmic cycles, then the methods presented here would be likely to identify it as an oscillation. The difference between sustained oscillations and brief bursts of band-limited activity has strong implications for the interpretation of these results [40]. To distinguish between oscillations and non-oscillatory bursts, future studies could extend the range of the behavioral time-series to test whether it is sustained over time. If future studies uncover significant behavioral oscillations using the AR surrogate or robust est. methods, they will need to design studies that are capable of disentangling ongoing oscillations from ‘bursty’ evoked patterns.

### Aperiodic dynamics in brain and behavior

Shuffling in time leads to spectral peaks that could reflect either periodic or aperiodic regularities in behavior. This finding does not invalidate the literature on attentional switching. Instead, it encourages us to consider these results in the context of the rich aperiodic temporal structure in perception and cognition. On short time-scales, for example, attention is impaired when two target events appear within around 100-500 ms of each other; this is called the ‘attentional blink’ [41, 42]. Furthermore, when people search through a visual scene, attention shows ‘inhibition of return’, with a suppression of perceptual processing of objects that have recently been attended [43]. Inhibition of return appears as quickly as 50 ms after a cue [44], and its effects can last for up to 3 s [45]. At longer time-scales (seconds to hours), behavior is correlated with itself over time, roughly following a 1/*f* spectrum. This 1/*f* pattern appears in reaction times, accuracy, and many other aspects of behavior [35, 36, 46].

Aperiodic dynamics are pervasive in neural recordings as well as in behavior [47], and a number of different mathematical approaches have been developed to investigate non-oscillatory features of neural time-series [48, 49]. The 1/*f* slope of a neural power spectrum correlates with the excitation/inhibition balance [50], and predicts a range of different behavioral variables [49, 51]. Aperiodic dynamics also appear in phenomena that are widely considered to be inherently oscillatory. For example, the power of high frequency oscillations often depends on the phase of lower-frequency oscillations [52]. A similar pattern emerges in non-oscillatory scale-free activity: the power of high frequency neural activity depends on the ‘phase’ of lower-frequency aperiodic activity [53]. For a second example, when spiking accompanies a consistent neural oscillation, the information carried by an action potential can depend on the oscillatory phase at which that action potential occurs [54]. This phenomenon is not limited to consistent neural oscillations - the information carried by a spike can also depend on the ‘phase’ of aperiodic low-frequency activity [55]. This type of non-oscillatory phase coding has been observed in behaving animals. Bats do not show any low-frequency oscillations in the hippocampal formation [56], but despite this lack of oscillations, hippocampal spiking locks to broadband fluctuations in the local field potential, and spike timing shows non-oscillatory ‘phase procession’ as the animal moves through space [57]. The findings summarized above suggest that non-oscillatory dynamics may play an important role in generating temporal structure in the brain and behavior [58–61], complementing the well-documented roles of neural oscillations [62].

### Characterizing aperiodic structure in behavior

How can we measure and conceptualize the aperiodic structure in behavior? In both the AR surrogate and the robust est. methods, aperiodic structure is measured by estimating the AR(1) coefficient that best fits the data. This AR(1) coefficient describes how strongly the signal correlates with itself across time, and therefore provides a measure of how quickly and consistently attention switches locations. Both of these methods could also be adapted to examine ‘power-law’ structure in behavior by fitting the data to a 1/*f^β^* spectrum instead of to an AR(1) process. This power-law approach has been widely applied to behavior on the scale of minutes to hours [35, 36, 46], but not yet on the scale of milliseconds to seconds. Furthermore, future research on attentional switching could examine concrete, non-repeating features of the behavioral time-series. This approach could identify features analogous to the attentional blink or inhibition of return, clarifying how attention dynamically samples the perceptual scene after a cuing event. An aperiodic view of temporal structure will enrich our understanding of attention and the neural processes that support it.

### Neural mechanisms of attentional dynamics

What neural mechanisms influence spontaneous attentional switching? Two major categories of models aim to explain rhythms in behavior via rhythms in brain activity. First, some researchers propose that a ‘pacemaker’ oscillation determines when attention switches between stimuli: perhaps endogenous oscillations at 3-8 Hz coordinate activity within a thalamo-cortical network, with one phase of each cycle facilitating sensory processing and another phase facilitating shifts in the attentional focus [5, 6]. In a second category of oscillatory models, some researchers propose that attentional switching may depend on competitive inhibition within visual cortex [4]. These competitive dynamics have been suggested to arise due to inter-hemispheric connections [63], or due to center-surround inhibition in overlapping receptive fields [17, 20].

The results reported here, however, raise the possibility that attentional switching could show non-oscillatory dynamics. Importantly, non-oscillatory patterns in behavior cannot be accounted for by a pacemaker oscillation or by mutual inhibition between overlapping receptive fields. What neural mechanisms could give rise to aperiodic attentional switching?

Aperiodic patterns of neural activity appear in a range of different behaviors and brain areas. For example, zebra finches sing motifs that are very similar across repetitions, and each repetition is accompanied by a highly stereotyped sequence of neural activity in the song-production area HVC [64]. Furthermore, rats show consistent aperiodic sequences of hippocampal spiking during working memory. These sequences are specific to the item being remembered [65]. Aperiodic patterns such as these may arise due to inherent properties of networks in the brain [58, 66–68], such as recurrent connections and a variety of time-dependent physiological processes (e.g. short-term synaptic plasticity) [69].

Research with artificial neural networks supports the idea that inherent network properties can give rise to aperiodic patterns of activity [58, 61]. In particular, sparsely-connected recurrent neural networks (RNNs) capture a number of the characteristics of attentional switching. For example, they show chaotic spontaneous dynamics [70], mirroring the way attention moves even without changes to the stimulus. Sparse RNNs are not limited to chaotic activity. These networks show reproducible dynamic patterns when they are stimulated with the same input [71], analogous to behavioral patterns like the attentional blink. Furthermore, a single RNN can flexibly generate multiple distinct behaviors, with different inputs corresponding to different dynamic responses [72, 73]. Correspondingly, different cues lead to different patterns of attentional switching (e.g. cue in the right versus left visual hemifield, [7]).

These computational and experimental findings motivate a novel hypothesis about the dynamics of attention: Non-oscillatory endogenous attentional switching may arise due to recurrent connections and time-dependent physiological processes within the attentional network. Prior research shows that the attentional focus moves as a function of the constantly-evolving population activity in brain areas such as the frontal eye fields (FEF) [24, 74, 75]. Reproducible patterns of aperiodic switching may emerge as a consequence of recurrent connections and time-dependent physiological processes in attentional areas in the brain. If this hypothesis is true, endogenous oscillations may not be necessary to account for patterns in attentional switching.

To discriminate between these oscillatory and non-oscillatory hypotheses, future studies could examine attentional switching in response to a sequence of multiple cue events. If attentional switching depends on ongoing oscillations that are reset by transient events (such as the cue stimulus) [7, 8, 76, 77], then the time-course of attention should not depend on whether the cue was preceded by another event. In contrast, recurrent network dynamics are sensitive to the initial conditions at the time that an event occurs [61, 69]. As a consequence of this context-dependence, this recurrent-connection hypothesis predicts that the time-course of attention will differ based on the amount of time between the two cue stimuli.

### Rhythms in perceptual sensitivity

Attentional switching is not the only aspect of perception that has been proposed to oscillate. A related literature shows robust evidence for rhythmic fluctuations in perceptual sensitivity [3]. Oscillations in perceptual sensitivity have been widely reported in behavioral studies (e.g. [78–80]). Electrophysiological studies show that perceptual sensitivity depends on the phase of ongoing neural oscillations [19, 81–92], (but see [93]). These studies of rhythms in sensitivity do not test for significant oscillations by shuffling the data in time, and are therefore not subject to the same statistical issues as the studies of attentional switching.

### Conclusions

The simulations presented here encourage us to reconsider temporal structure in attentional switching. Attention may move along with endogenous neural oscillations; alternatively, attentional switching may depend on aperiodic patterns that arise from recurrent connections and time-dependent physiological processes. In this article, I provide tools to distinguish between oscillatory and non-oscillatory structure in brief, noisy time-series. Furthermore, I demonstrate that prior studies show no evidence for behavioral oscillations when they are re-analyzed with these novel tools. To understand the neural systems supporting attention, future studies should explicitly test for periodic versus aperiodic temporal structure in behavior.

## Methods

In this study, computational simulations of behavioral experiments were analyzed according to standard procedures from highly-cited papers. The simulations and analyses followed the details from two prominent studies: Landau and Fries (2012) [abbreviated to LF2012], and Fiebelkorn, Saalmann, and Kastner (2013) [abbreviated to FSK2013]..

All p-values reflect two-tailed tests except where otherwise noted.

### Simulated behavioral experiments

Behavioral studies were simulated using the details of the experiments in LF2012 and FSK2013. In both experiments, participants were first presented with visual stimuli on the screen. After a short delay, participants saw a cue stimulus intended to attract spatial attention and reset ongoing cortical dynamics. After a variable delay, a faint target stimulus appeared briefly at either the cued location or an uncued location. These studies then considered changes in accuracy as a function of the delay between the cue and target.

In these simulations, each experiment began with an idealized accuracy time-course. This is the timecourse of accuracy that would be obtained after running an infinite number of trials. For each trial, a cue-target delay is randomly selected, with a balanced number of trials at each delay. Accuracy for each trial was randomly determined as a function of the idealized accuracy time-course. For example, if a trial is selected for a cue-target delay of 0.5 s, and the idealized accuracy at 0.5 s is 60%, then that trial has a 60% chance of being a hit and a 40% chance of being a miss.

#### Experiment details: LF2012

For simulations following LF2012, time-courses were simulated with cue-target delays of 0.15 s to 1.0 s, sampled at 60 Hz. I simulated data from 16 subjects, with each subject having 104 trials at each location.

#### Experiment details: FSK2013

For the simulations following FSK2013, time-courses were simulated with cue-target delays of 0.3 to 1.1 s, sampled at 60 Hz. I simulated data from 15 subjects, with each subject having 441 trials at each location.

#### Experiment details: AR surrogate and Robust estimate analysis

For the simulations using the other two methods (AR surrogate and robust est.), experiments were simulated as in LF2012.

### Identifying rhythms in simulated behavior

We simulated behavioral experiments following the analysis pipelines from LF2012 and FSK2013. In addition, experiments were simulated using a novel permutation method that uses autoregressive models to generate a surrogate distribution. Finally, experiments were simulated using a method for determining spectral peaks in climatic time series (Mann and Lees, 1996; abbreviated to ML1996). All analyses only considered frequencies below 15 Hz (12 Hz in the FSK2013 analysis, reflecting the plotted data).

#### Shuffling in time: LF2012

For this analysis method, we search for rhythms in behavior following LF2012. To derive the spectrum, accuracy is first averaged over trials and subjects at every cue-target delay. The data are then detrended with a 2nd-order polynomial, tapered with a Hanning window, and padded to 256 samples. Finally, the amplitude spectrum is obtained by taking the magnitude of a discrete Fourier transform (DFT) of this time-series.

A randomization procedure is used to determine the statistical significance of peaks in the spectrum. For *k* = 500 permutations, the time of the cue-target delay is randomly shuffled among all the trials, and the spectrum is calculated following the same procedure as used in the empirical data. P-values in each simulated experiment are computed as the proportion of values at each frequency that have greater spectral magnitude than the empirical spectrum, followed by Bonferroni correction for multiple comparisons across frequencies.

#### Shuffling in time: FSK2013

This analysis method follows the procedures in FSK2013. To derive the spectrum, accuracy is first averaged over trials and subjects in overlapping time-windows with width 0.05 s, advancing by steps of 0.01 s. This accuracy time-series is then detrended with a 2nd-order polynomial, tapered with a Hanning window, and padded to 128 samples. Finally, the spectrum is computed by taking the magnitude of the DFT of this time-series.

Statistical significance is determined as in LF2012, by shuffling the trials in time and recomputing the spectra (*k* = 1000), correcting over multiple comparisons across frequencies using the false discovery rate [94].

#### Robust estimation of background noise: ML1996

In addition to the randomization analyses popular in cognitive neuroscience, I used a technique that is common in geology and climate science [33]. This procedure was developed to identify rhythms in geological time-series, and to isolate these rhythms from a background of autocorrelated noise. We apply this analysis to the time-course of average accuracy at each cue-target delay. The mean of the time-course is subtracted, and the spectrum is computed using Thomson’s (1982) multi-taper procedure [95]. For appropriate spectral smoothing, we selected a time-bandwidth parameter of 1.5 with 2 tapers. We then estimate the aperiodic background spectrum by smoothing the multi-taper spectrum with a median filter (width = 7), and using this robustly-smoothed spectrum to fit an estimate of an AR(1) spectrum approximating the aperiodic background activity:

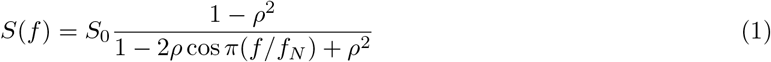

where *f* is frequency, *f_N_* is the Nyquist frequency, *S*_0_ is the average value of the power spectrum, and *ρ* is the AR(1) coefficient. Finally, we test for statistically significant periodic components by taking the ratio of the multi-taper amplitude spectrum against the robust estimate of the AR(1) background spectrum, separately for each frequency, and comparing this to a χ^2^ distribution with degrees of freedom equal to 2 × (number of tapers). For further details, see [33].

#### AR surrogate

Data were also analyzed using a novel method to test for significant oscillations in autocorrelated time-series. This method uses AR(1) models to generate a surrogate distribution for non-parametric randomization tests. First, accuracy is averaged at each cue-target delay. We then estimate an autoregressive model with 1 parameter for this time-series. An AR(1) model captures the aperiodic temporal structure in the behavioral time-series, but cannot generate consistent oscillations. This fitted AR(1) model is then used to generate a surrogate distribution of time-courses (*k* = 2000). The time-courses generated by this model preserve the aperiodic structure of the empirical data, but lack any periodic components. The time-courses from the empirical data and the AR-generated surrogate signals are then de-meaned, and the spectra are computed by taking the magnitude of the DFT. No tapering or zero-padding was applied. In preliminary analyses, tapering was found to drastically reduce both the power of this analysis and the accuracy of the frequency estimates. We correct for multiple comparisons across frequencies using a cluster-based permutation test [34] (cluster threshold alpha: 0.05; cluster statistic: summed z-score).

### Simulated behavioral time-courses

To determine how the different analysis methods respond to different types of temporal structure in behavior, I simulated literatures of 1000 experiments for each combination of analysis method and temporal structure. Each simulated experiment began with an idealized accuracy time-course that was the same across all participants and trials within that experiment. This idealized accuracy formalizes the temporal structure of attentional switching after the cue stimulus. For each trial, a cue-target delay was randomly selected. We then randomly determined whether that trial was a hit or a miss based on the idealized accuracy for that cue-target delay. For example, if one simulation has an idealized accuracy of 70% at a cue-target delay of 0.60 s, then each trial for that time-point is determined as a weighted coin-toss, with *P*(hit) = 0.7 and *P*(miss) = 1 - 0.7 = 0.3.

We tested four types of aperiodic temporal structure. ‘Fully random’ simulations contained no temporal structure at all, with no consistency over trials. These time-series were simulated with an idealized accuracy time-course with *P*(hit) = 0.5 at every cue-target delay. For ‘white noise’ simulations, idealized accuracy was generated by sampling a random Gaussian process. ‘Random walk’ simulations were generated with a Gaussian random walk; this is equivalent to a power-law spectrum with an exponent of 2 (1/*f*^2^). ‘AR(1) noise’ simulations were generated with a 1-coefficient Gaussian autoregressive process with *β* = 0.5. For simulations that included consistency across trials (all except ‘fully random’), the idealized accuracy was rescaled to approximate the accuracy range in the behavioral literature: [0.5, 0.7].

Next, I tested how the different analyses reconstruct true oscillations in behavior. For these simulations, the idealized accuracy time-course was generated as a sine-wave with randomized phase, at frequencies from 2–12 Hz in steps of 1 Hz. Amplitude of these behavioral oscillations (corresponding to the range between minimum and maximum accuracy) was varied from 0.1–0.6 in steps of 0.1. Mean accuracy was held at 0.5. These oscillations were then added to a random walk generated as described above, and the resulting time-series was used as an idealized accuracy time-course.

To measure how precisely each analysis recovers true oscillations, I computed the error in recovered rhythms: (the frequency of significant components) - (the frequency of the simulated oscillation). The median of the absolute value of the error (median absolute error) was computed separately for each frequency and amplitude of simulated oscillations. Linear regressions were then used to test for differences between analysis methods. The median absolute error was modeled as a function of the analysis method, with terms to control for the frequency and amplitude of the simulated oscillation. These regressions included one observation for each combination of simulated frequency and amplitude. I report the regression coefficient reflecting the analysis method, along with 95% confidence intervals, t-statistics, and p-values.

To quantify the selectivity of each analysis method, I computed a detection ratio: (the rate of correct positive results when an oscillation is present) / (the rate of false positive results when no oscillation is present). Linear regressions were used to test for differences between methods, as in the analysis of the median absolute error.

### Re-analysis of publicly-available data

Data from 4 previous studies was retrieved from public repositories [15, 18, 22, 26]. Every test using shufflingin-time was re-analyzed using the two novel alternative methods. For each test, I reproduced the aggregated behavioral time-course, and verified it against the plot of the time-course in the original paper. These timeseries were then re-analyzed using the AR surrogate and robust est. methods. I counted the number of statistically significant results (*p* < .05 after correcting for multiple comparisons) reported in the published paper, and compared this with the count of statistically significant results from the alternative methods.

## Data availability

Results of all simulations will be made available in an OSF repository upon puplication. The re-analyzed data are available at the following publicly-available repositories.

- Ho et al (2017) [15]: https://ars.els-cdn.com/content/image/1-s2.0-S0960982217313209-mmc2.xlsx
- Davidson et al (2018) [18]: https://figshare.com/projects/Crossmodal_binocular_rivalry_attention_sampling_project/56252
- Senoussi et al (2019) [22]: https://osf.io/2d9sc/?view_only=6ef3f85d9f944d27b23fc7af5a26f087
- Michel et al (2021) [26]: https://osf.io/de4bu/

## Code availability

All code needed to perform the analyses and generate the plots will be made available at a publicly-accessible repository upon acceptance of the paper.

## Notes

### Competing Interest Statement

The authors have declared no competing interest.

